# Structural and Functional Consequences of Age-Related Isomerization in α-Crystallins

**DOI:** 10.1101/364497

**Authors:** Yana A. Lyon, Dylan L. Riggs, Miranda P. Collier, Matteo T. Degiacomi, Justin L.P. Benesch, Ryan R. Julian

**Affiliations:** Department of Chemistry, University of California, Riverside, 501 Big Springs Road, Riverside, CA 92521, USA; Department of Chemistry, Physical and Theoretical Chemistry Laboratory, University of Oxford, South Parks Road, Oxford OX1 3QZ, UK; Department of Chemistry, Durham University, South Road, Durham DH1 3LE, UK

**Keywords:** lens, mass spectrometry, radical, photodissociation, epimer, isomer, long-lived protein

## Abstract

Long-lived proteins are subject to spontaneous degradation and may accumulate a range of modifications over time, including subtle alterations such as isomerization. Recently, tandem-mass spectrometry approaches have enabled the identification and detailed characterization of such peptide isomers, including those differing only in chirality. However, the structural and functional consequences of these perturbations remain largely unexplored. Here we examine the site-specific impact of isomerization of aspartic acid and epimerization of serine in human αA- and αB-crystallin. From a total of 81 sites of modification identified in aged eye lenses, four (^αB^Ser59, ^αA^Ser162, ^αB^Asp62, ^αB^Asp109) map to crucial oligomeric interfaces. To characterize the effect of isomerization on quaternary assembly, molecular dynamics calculations and native mass spectrometry experiments were performed on recombinant forms of αA- and αB-crystallin that incorporate, or mimic, isomerized residues. In all cases, oligomerization is significantly affected, with epimerization of a single serine residue (^αA^Ser162) sufficing to weaken inter-subunit binding dramatically. Furthermore, phosphorylation of ^αB^Ser59, known to play an important regulatory role in oligomerization, is severely inhibited by serine epimerization and altered by isomerization of nearby ^αB^Asp62. Similarly, isomerization of ^αB^Asp109 disrupts a vital salt-bridge with ^αB^Arg120, a loss previously shown to yield aberrant oligomerization and aggregation in several disease variants. Our results illustrate how isomerization of amino-acid residues, which may seem like a minor structural perturbation, can have profound consequences on protein assembly and activity by disrupting specific hydrogen bonds and salt bridges.

**Significance Statement:** Proteins play numerous critical roles in our bodies but suffer damage with increasing age. For example, isomerization is a spontaneous post-translational modification that alters the three-dimensional connectivity of an amino acid, yet remains invisible to traditional proteomic experiments. Herein, radical-based fragmentation was used for isomer identification while molecular dynamics and native mass spectrometry were utilized to assess structural consequences. The results demonstrate that isomerization disrupts both oligomeric assembly and phosphorylation in the α-crystallins, which are long-lived proteins in the lens of the eye. The loss of function associated with these modifications is likely connected to age-related diseases such as cataract and neurodegenerative disorders, while the methodologies we present represent a framework for structure-function studies on other isomerized proteins.

## Introduction

Long-lived proteins are important but often underappreciated, with recent findings illustrating their pervasiveness within critical organs and suggesting that their chemistry and biology should not be ignored.^1^ Longevity renders proteins susceptible to degradation and accumulation of post-translational modifications (PTMs). Among these modifications are non-enzymatic, spontaneous changes including truncation, cross-linking, oxidation, deamidation, isomerization, and epimerization.^1^ Aspartic acid residues are most prone to isomerization,^2,3^ readily forming a succinimide ring following attack of the sidechain by the peptide backbone. The succinimide is susceptible to racemization and can reopen in two ways, ultimately yielding four isomers: L-Asp, D-Asp, L-isoAsp, and D-isoAsp.^4^ After deamidation, asparagine can also yield four isomers of aspartic acid, but this process involves chemical modification in addition to isomerization. Serine is also frequently found to undergo isomerization in long-lived proteins, epimerizing to D-Ser. Isomerization is difficult to detect because it does not lead to a change in mass, and is consequently invisible to mass spectrometry (MS)-based methods typically employed during proteomic analyses.^5,6^ As a result, isomerization is not widely studied, likely frequently overlooked, and its consequences are largely unknown.^7^

The crystallin proteins of the eye lens, in which there is no protein turnover, are among the longest-lived proteins in the body.^8^ They are ideal targets for studying spontaneous degradation pathways that occur due to aging. The most abundant crystallins in humans, αA and αB, are important molecular chaperone^9^ and regulatory proteins.^10^ While αA is localized almost exclusively to the eye lens,^11^ αB is found throughout the body.^12,13^ Crystallin malfunction due to mutation or accumulation of PTMs is associated with a variety of diseases including cataract, cardiomyopathies, motor neuropathies, and neurodegeneration.^14-16^

αA and αB are members of the small heat-shock protein (sHSP) family,^9^ with structures characterized by a highly conserved “α-crystallin domain” (ACD),^17^ flanked in both proteins by less ordered N- and C-terminal regions (Fig. 1A). Outside of the ACD, αA and αB share modest sequence homology,^11^ and in both cases structural details have proven hard to come by. αA and αB use a number of interfaces to self- and co-assemble into large, polydisperse and dynamic oligomers. The ACD mediates dimerization, and the dimers associate into oligomers via an interface between a palindromic sequence in the C-terminal region of one dimer and the ACD of another, as well as interactions involving N-terminal regions (Fig 1B).^18-20^ Perturbation of the dimer interface, as occurs in the well-known R120G variant of αB,^21^ or mutations in the terminal regions, can lead to protein aggregation and malfunction.^22^ The function and localization of αB is also partially regulated by phosphorylation of three serines in the N-terminal region, with dysregulation leading to disease.^23-27^ These data are consistent with modification of interfacial residues leading to aberrant assembly and compromised function in the cell.

**Fig. 1.**
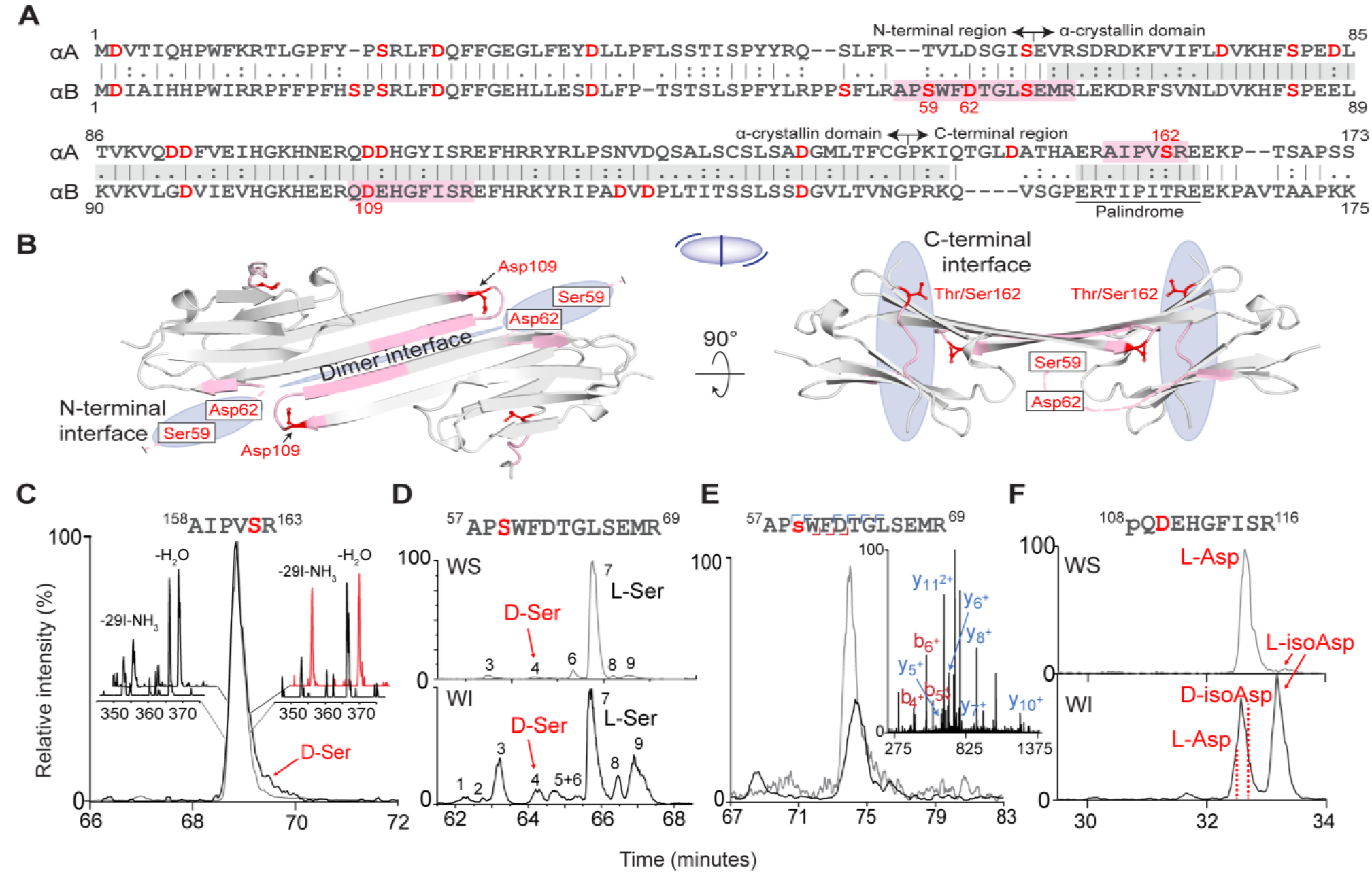
(A) Sequence alignment of human αA and αB with designated boundaries between domains. Red residues indicate Asp/Ser sites within peptides from 72-year-old human lens identified in numerous isomeric forms by RDD-MS. Pink regions indicate isomer-containing peptides mapping to oligomeric interfacial regions. (B) Partial crystal structure of α-crystallin (αB, PDB: 4M5S) comprising residues shaded grey in (*A*). Consequences of isomerization at red labeled residues were investigated in the present study. Blue shaded regions indicate oligomeric interfaces. In the middle, a schematic representation of the crystal structure with blue lines indicating the inter-dimer interface (center) and C-terminal interactions with other monomers (peripheral). On the right, rotated top view of C-terminal binding to the α-crystallin domain. The structure of αB is used for illustration of isomerized regions for both proteins, which intermix and share high structural similarity. (C) Extracted ion chromatogram of 4IB-AIPVSR (where 4IB = *para*-iodobenzoate modification) from the WI (black) and WS (grey) fractions of the cortex. Insets: RDD mass spectra from the front and back of each peak. Comparing the relative fragment intensities of m/z 350.25 and 364.00 for the WI fraction yields an R_isomer_ score of 2.7, confirming the presence of two isomers. (D) Extracted ion chromatogram of ^57^APSWFDTGLSEMR^69^ from the WI/WS nucleus, revealing abundant isomerization in the WI fraction including D-Ser. (E) Extracted ion chromatogram of phosphorylated ^57^APsWFDTGLSEMR^69^ detected in the WS cortex (grey) and WI cortex (black) revealing far less isomerization, where s denotes site of serine phosphorylation. Inset: collision-induced dissociation of the phosphorylated peptide in the WS fraction with diagnostic peaks that pinpoint the site of phosphorylation to Ser59. (F) Extracted ion chromatograms from the WS and WI fractions of ^108^pQDEHGFISR^116^ from the cortex (pQ=pyroglutamate). In the WS fraction, the major peak contains L-Asp and the minor peak is L-isoAsp. In the WI fraction, the peak at ∼33 minutes contains both L-Asp and D-isoAsp; and the final peak is L-isoAsp.

We have recently detected 81 sites of isomerization in human αA and αB isolated from the eye lenses of aged donors^28^ by using high-throughput radical-directed dissociation RDD-MS experiments.^2^ Here we have identified four age-related isomerization sites in αA and αB (^αB^Ser59, ^αA^Ser162, ^αB^Asp62, ^αB^Asp109) that reside in regions critical for oligomerization, and demonstrate how key structural interactions are disrupted by these modifications. Native MS^29,30^ reveals that the interface between the C-terminal region and the ACD is dramatically weakened upon conversion of L- to D-Ser. Epimerization abrogates serine phosphorylation, with isomerization of neighboring residues also interfering with kinase recognition. Molecular dynamics (MD) simulations show that isomerization of an aspartic acid at the dimer interface leads to cleavage of a salt-bridge, and precipitates the aberrant oligomerization observed in native MS data. Our results demonstrate that age-related, isobaric modifications of individual amino-acid residues can have significant impacts on the quaternary structure and function of α-crystallins, and suggest that similar “invisible” PTMs in other long-lived proteins may have an important influence on age-related diseases.

## Results and Discussion

### αA and αB accumulate isobaric PTMs at their interfaces

The sequence alignment and relevant structural regions for αA and αB crystallin are shown in Fig. 1A. RDD-MS^2^ was used to identify potential sites of isomerization in αA and αB, and illustrative results are shown for the nucleus and cortex of a 72-year-old human lens, although similar results were obtained from lenses of other ages.^28^ These experiments revealed isomerization in peptides spanning numerous sites, including many with the potential to disrupt key structural interactions (Fig. 1B). For additional details on isomer identification, see supporting information Figure S1. We will focus on three regions known to be involved in oligomerization that are found within the tryptic peptides: αA-^158^AIPVSR^163^, which lies in the part of the C-terminal region responsible for assembly; αB-^57^APSWFDTGLSEMR^69^, which encompasses residues involved in N-terminal region interactions; and αB-^108^QDEHGFISR^116^ which lies in the ACD and spans a large portion of the dimer interface.

Performing RDD-MS across the chromatographic peak for αA-^158^AIPVSR^163^ reveals product ions at 350.25 m/z and 364.00 m/z, whose ratio varies with elution time for the water-insoluble (WI) fraction, as illustrated by the snapshots in Fig. 1C. The RDD spectra for the water-soluble (WS) fraction do not vary as a function of retention time. The changing ratios for the WI fraction are meaningful because in RDD experiments,^31^ a radical is photolytically created at the same, atomically precise location in both isomers. Subsequent collisional activation stimulates migration of the radical and fragmentation of the peptide. Differences in three-dimensional structure due to isomerization lead to differences in the abundance of certain fragmentation channels. Therefore, the varying product ion ratios for the WI fraction in Fig. 1C reveal epimerization at ^αA^Ser162. Further analysis^32^ enables quantification of the abundance of ^αA^D-Ser162 at 8% (see Fig. S3 for details). Similarly, the likeness of the RDD spectra for the WS fraction indicates absence of D-Ser. This approach can also be applied to more complex systems, as illustrated by the chromatograms for separation of αB-^57^APSWFDTGLSEMR^69^, from which nine different isomers were identified in the WI fraction (Fig. 1D). Synthetic standards of selected candidates were used to determine that the original all-L peptide comprises only 14% of the total abundance in the WI fraction (peak 7, which dominates the WS fraction), while the ^αB^D-Ser59 epimer is present in 5.6% abundance (peak 4) in the WI fraction and 1.4% in the WS fraction. Interestingly, when αB-^57^APSWFDTGLSEMR^69^ is phosphorylated at ^αB^Ser59, only a single isomer is detected in the WS fraction (Fig. 1E) and a small number of isomers are detected in the WI fraction. The WS and WI fractions of αB-^108^QDEHGFISR^116^ both contain multiple isomers of ^αB^Asp109 (Fig. 1E), but the relative abundance of ^αB^L-isoAsp109 increases from 5.1% to 55.4% in the WI portion, suggestive of structural consequences of this PTM. These data demonstrate that isobaric PTMs are prevalent in αA and αB, including modifications found at interfacial regions responsible for assembly into the functional oligomeric forms that appear to influence solubility.

### Epimerization compromises the interface between the ACD and C-terminal region

^αA^Ser162 is an epimerized residue located adjacent to the highly conserved IXI/V motif in the palindrome of the C-terminal region (Fig. 1A-C). This part of the sequence enables the flexible C-terminal regions to form bi-directional domain-swap interactions between dimers, by binding the groove between β4 and β8 on an adjacent ACDs. ^33^ These interactions are crucial for proper oligomerization, and point mutations can influence the kinetics and thermodynamics of assembly.^34^ Examination of the available crystal structures of αA (Fig. 2A, truncated variants from *Bos taurus* and *Danio rerio*, PDB 3L1F and 3N3E) reveals that the C-terminal region runs across the groove in the β8 → β4 direction and the inverse direction (β4 → β8), such that residue 162 (Ser for αA or Thr for αB) is involved in a hydrogen bond with the backbone carbonyl of ^αA^Val87 or ^αA^Cys131, respectively. 200 ns MD simulations reveal these interactions to be formed ∼5% of the time.

**Fig. 2.**
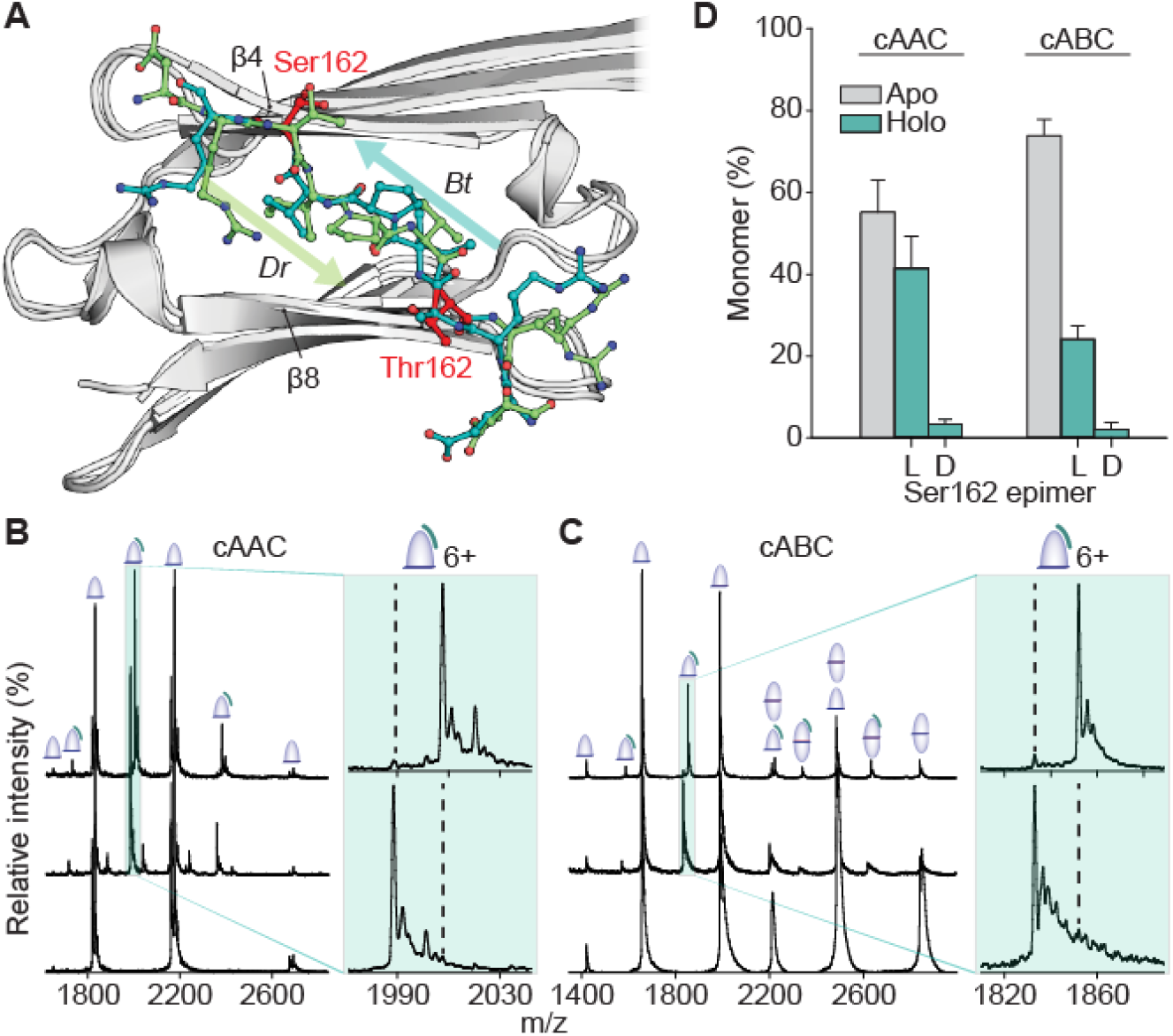
Binding competition experiments reveal a strong preference for L- over D-Ser162 within both α-Crystallin oligomeric interfaces. (A) Aligned crystal structures of the ACD (grey) and C-terminal residues 156-164 of bovine (*Bt*) (teal) and zebrafish (*Dr*) (green) αA-crystallin. Arrows indicate orientation (N → C) of C-terminal binding. Potential isomer Ser162 (alternate conformations) and Thr162 in the equivalent position are shown in red. (B) Native mass spectra of cAAC alone (bottom, 10 μM) and mixed with 40 μM C-terminal peptides ERAIPVSRE (Ct) and GERAIPVsREG (G-Ct-G) with incorporation of L-Ser162 and D-Ser162, respectively (middle) and with peptides of reversed Ser162 chirality (top). Magnified region (green) shows the 6+ charge state of cAAC-peptide complexes evidencing preferential binding to L-epimer peptides. Dotted lines indicate expected positions of D-epimer-bound peaks. (C) Identical experiment using cABC. Whereas dimer was only present in trace amounts for cAAC and, therefore, excluded from panel (A), cABC dimer and peptide-bound dimer peaks are indicated. Magnified: 6+ charge state of cABC-peptide complexes. (D) Relative fractions of monomer bound to no peptides, L-peptides, or D-peptides in competition experiments, using all 5+, 6+ and 7+ charge state intensities for quantitation. Error bars represent 95% confidence intervals.

To quantify the effect of ^αB^Ser162 epimerization, we conducted a series of native MS experiments involving the ACDs of αA and αB, and isomeric variations of the peptide ERAIPVSRE, the palindromic part of the C-terminal tail of αA. The core domains (cAAC = core αA-crystallin, cABC = core αB-crystallin), when analyzed alone, returned charge-state series corresponding to a monomer (αA), and both monomer and dimer (αB) (Fig. 2B-C, lower spectra). This is suggestive of the αA dimer interface being slightly weaker than that of αB. Competition binding experiments between the ACD and palindromic peptides containing either D- or L-Ser162 are shown in Fig. 2B-C, middle spectra. The canonical all L-amino acid form of the peptide, ERAIPVSRE, and a D-isomer variant made heavy by addition of glycine to both termini, GERAIPVsREG (s = D-Ser), were added at a 4:1 peptide/ACD molar ratio. Peptide binding to both αA and αB ACDs was observed, consistent with nuclear magnetic resonance spectroscopy (NMR) data,^35^ and previous observations that αA and αB freely intermix to form higher order oligomers.^19^ The dominant (>95%) adduct observed corresponds to the binding of the canonical peptide (see insets, lower spectra). Furthermore, the abundance of ACD dimers decreases for αB (Fig. 2C), consistent with allosteric interference in dimer binding, as observed previously.^34^

To eliminate the possibility that the additional glycine residues might be responsible for the difference in binding, the inverse experiment was conducted with the heavy L-isomer for both ACDs (Fig. 2B-C, and insets, upper spectra). Binding of the L-isomer dominates in both systems (Fig. 2D), and the data can be used for quantification^36^ of the dissociation constants: L-Ser:cAAC = 48 µM, L-Ser:cABC = 115 µM, D-Ser:cAAC = 650 µM, D-Ser:cABC = 1380 µM. Based on these values, epimerization leads to destabilization of ΔΔG = ∼6 kJ/mol. All of these results are consistent with D-^αA^Ser162 significantly inhibiting proper interaction between the C-terminal palindrome and the β4-β8 groove in the ACD.

We have previously shown that removal of the side-chain at the equivalent position in αB (^αB^Thr162Ala) leads to a weaker interaction in the peptide-ACD system and faster subunit-exchange of full-length αB.^34^ The present data suggests that epimerization of ^αA^Ser162 will similarly affect the dynamics, and consequently chaperone activity, of αA. Furthermore, because the diversity of binding modes between the C-terminal tail and the ACD promotes polydispersity and aids in preventing crystallization,^33^ disruption of this binding may underlie the presence of the isomerized peptide in the WI lens fraction.^33^ Since both chaperone activity and polydispersity are likely crucial for lens transparency, ^22,37^ epimerization of ^αA^Ser162 may be a contributor to age-related protein aggregation within the lens.

### Phosphorylation is precluded by epimerization of ^αB^Ser59 and affected by isomerization of ^αB^Asp62

Ser59 is an important site that has previously been demonstrated to isomerize more abundantly in cataractous lenses.^38^ As shown in Fig. 1D, the peptide containing ^αB^Ser59 and ^αB^Asp62 is also highly isomerized in a normal lens. In addition, ^αB^Ser59 is a primary site of phosphorylation and helps regulate the oligomeric size and activity of αB,^39^ though phosphorylation is primarily observed in the younger cortex within the lens.^40^ Solid-state NMR data suggests that this residue is involved in inter-monomer contacts.^41^ To investigate the influence of isomerization of ^αB^Ser59 and ^αB^Asp62, we incubated the corresponding synthetic isomers of FLRAPSWFDTG-NH_2_ with the native kinase, MAPKAPK-2.^42^ The extracted ion chromatograms for both peptides reveal that the L-Ser59 isomer is the best substrate for the kinase (Fig. 3B). The ratio of L/D-Ser59 phosphorylation is ∼240/1 after 2 hours and ∼350/1 after 12 hours incubation. Interestingly, examination of the phosphorylated peptide in the lens reveals minor isomerization (Fig. 1E), offering sharp contrast to the abundant isomerization of the unmodified peptide. This suggests that modifications elsewhere on the peptide may also inhibit phosphorylation, or that phosphorylation prevents isomerization, or both. Additional *in vitro* experiments confirmed that phosphorylation is inhibited by D-isoAsp62, but slightly enhanced by D-Asp or L-isoAsp at the same position (Fig. 3B). In sum, the native kinase activity is affected by all three non-native Asp isomers. These results agree with previous experiments that failed to isolate phosphorylated D-Ser from erythrocytes.^43^

**Fig. 3.**
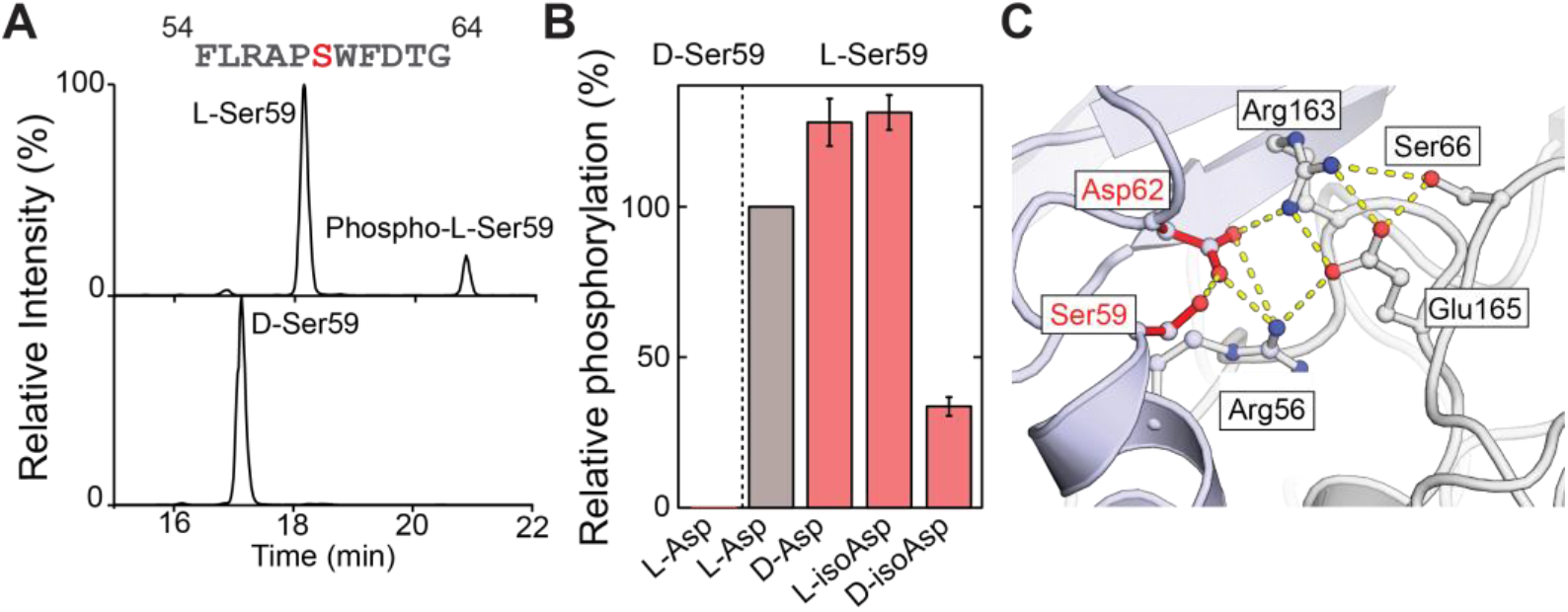
(A) Extracted ion chromatograms following incubation of L-Ser and D-Ser FLRAPSWFDTG-NH_2_ and FLRAPsWFDTG-NH_2_ with MAPKAPK-2 reveal that D-Ser is not a competitive phosphorylation substrate. (B) Relative degree of phosphorylation for Asp and Ser isomers of FLRAPSWFDTG-NH_2_. (C) Salt-bridge model (PDB 2YGD) of an N-terminal inter-oligomeric interface involving Ser59 and Asp62. Hydrogen bonds are shown using dashed yellow lines.

Ser59 and Asp62 have also been suggested to be involved in an interfacial salt-bridge cluster with Arg163 and Glu165 of an adjacent monomer.^44^ In this model, all four residues are in close proximity, with isomerization of Asp62 or Ser59 both likely to perturb the dynamics of the salt bridge and disrupt the interface (Fig. 3C). Similarly, phosphorylation of Ser59, which resides 2.4 Å (O-O distance) from Asp62, is likely to disrupt this salt bridge network and may account for the reduced oligomer size observed for phosphorylation mimics of Ser59.^45^

### Isomerization of ^αB^Asp109 breaks an interfacial salt-bridge and leads to insolubility

Figure 1F illustrates abundant isomerization of ^αB^Asp109, which is known to form an inter-monomer salt-bridge with ^αB^Arg120 in the AP_II_ register,^33,41,46^ the most populated state in solution.^47^ Mutation at either site in the salt bridge frequently leads to malfunction and disease.^46-49^ For example, the R120G mutant is genetically linked to desmin-related myopathy,^50^ while mutation of ^αB^Asp109 is associated with myofibrillar myopathy^51^ and cardiomyopathy.^52^ To probe the structural consequences of ^αB^Asp109 isomerization on the dimer interface of αB, *in silico* mutation and MD simulations were utilized, after parameterizing the force field for the isomeric amino acids.

The results obtained from all-atom simulations extending >150 ns are illustrated in Fig. 4. The Asp109-Arg120 salt bridge is stable for the L-Asp isomer, with a mean acceptor to donor hydrogen bond distance under 2 Å (Fig. 4A, left). For all other isomers, D-Asp109, L-isoAsp109, or D-isoAsp109, the salt-bridge with Arg120 is disrupted, and the average distances increase significantly (Fig. 4A, left). To determine the influence Asp isomerization has on the stability of the dimer interface, we monitored the distances between the final backbone hydrogen bond partners (His111 and Arg123). Reasonable hydrogen bond distances are only maintained for the L-Asp isomer, with all other isomers producing elongated distances (Fig. 4A, right). These results are illustrated by structural snapshots in Fig. 4B. For the L-Asp isomer (upper left), a backbone hydrogen bond between His111 and Arg123 links together the ends of β6+7 strands. For the other three isomers, these partners have been shifted to non-interacting distances due to disruption of the β-sheet dimer interface. The results from these simulations rationalize the observed partitioning of each isomer extracted from the lens (Fig. 1F). The abundance of isomerized Asp109 residues is much higher in the WI fraction, while L-Asp109 is the dominant isomer present in the WS fraction. This suggests that perturbation of Asp109 leads to enhanced protein aggregation, consistent with similar observations for the R120G mutation.^49^

**Fig. 4.**
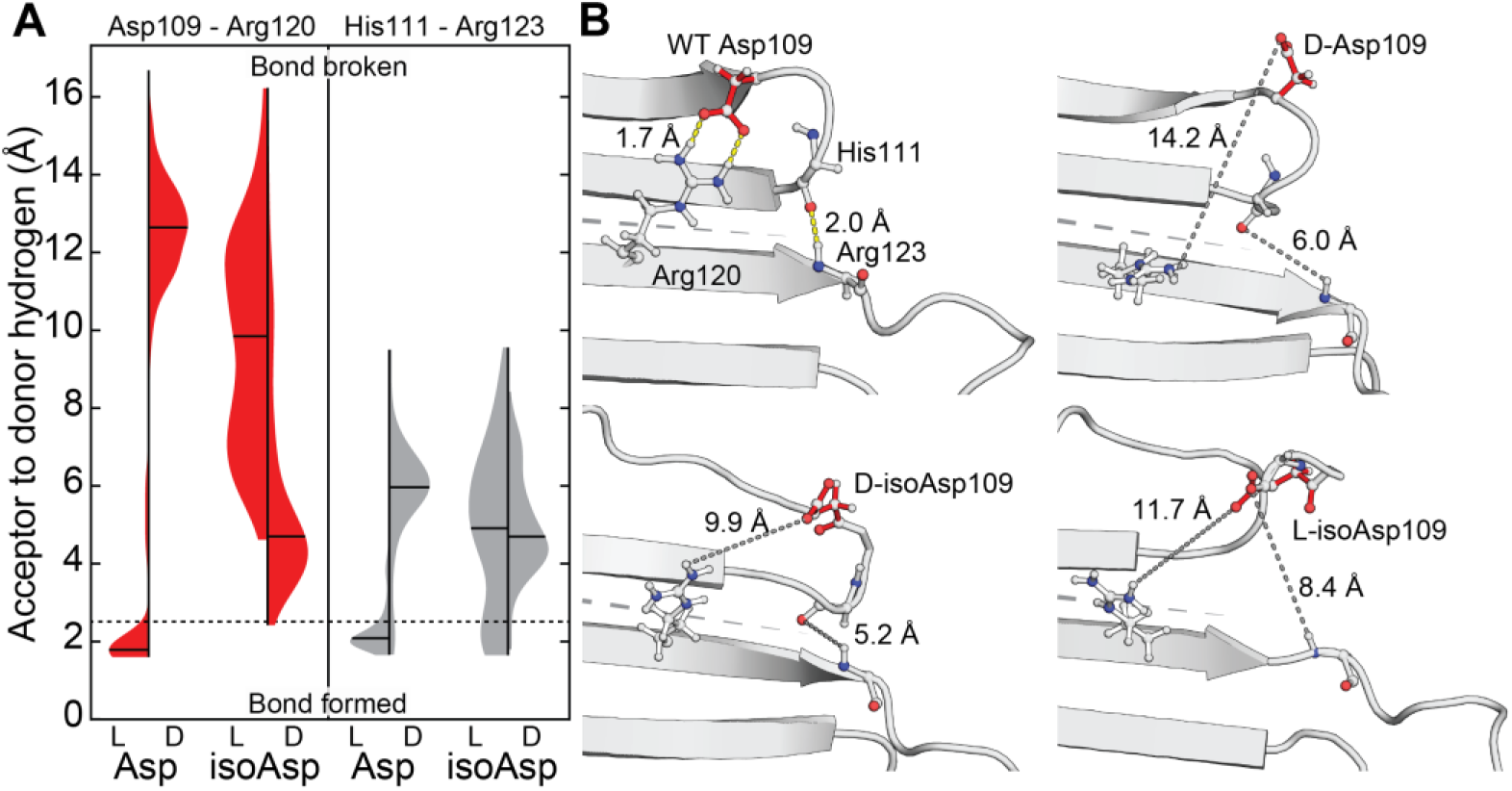
(A) Distance distributions between Asp109 and Arg120 (left,red), and His111 and Arg123 (right, grey) from MD simulations. Violin plots are shown for each isomer of Asp109; means are marked with black lines. Lengths <2.5 Å can be considered to correspond to bond formation (neglecting consideration of the bond angles), while those longer represent absence of the bond (boundary demarcated by dashed line). In all isomers other than L-Asp, the hydrogen-bond donor and acceptors are located too far apart for bond formation, the vast majority of the time. (B) Representative frames from MD simulations showing breakage of hydrogen bonds profiled in (A) and resultant interface destabilization. Yellow dashes indicate H-bonds; short gray dashes show concomitant distances following isomerization of Asp109; long gray dashes mark the antiparallel dimer interface. His111 and Arg123 side-chains have not been shown, for clarity.

### Mimicking breakage of interfacial bonds by isomerization leads to aberrant oligomerization

Our examination of the amino-acid environment around both Ser59 and Asp109 revealed that in both cases modification at these sites would be likely to impact oligomerization. Specifically, we noted that ^αB^Ser59 appears to be part of a network of salt bridges, into which a bulky negative charge seems unlikely to be accommodated without significant rearrangement. To test this prediction, we generated the phosphorylation mimic ^αB^Ser59Asp and compared it to the wild-type by using native MS. Both proteins gave mass spectra featuring a broad region of signal at high m/z, indicative of a polydisperse ensemble of oligomers, and consistent with previous spectra of αB (Fig. 5A).^53^ Notably, the signal is at slightly lower m/z values for the phosphomimic, consistent with a shift to smaller stoichiometries. In order to quantify this change, we performed collisional activation to remove highly charged monomers from the parent oligomers, resulting in lower ‘charge-stripped’ oligomers that are well resolved and can be deconvolved into an oligomeric distribution (Fig. 5B).^53^ Ser59Asp yields a distribution centered on an 18mer, smaller than the wild-type, and with a stronger preference for oligomers with an even number of subunits (Fig. 5C). This suggests that phosphorylation of Ser59, which regulates activity and localization of αB,^39^ leads to destabilization of the larger oligomers, likely at inter-dimer interfaces involving the N-terminal region.

**Fig. 5.**
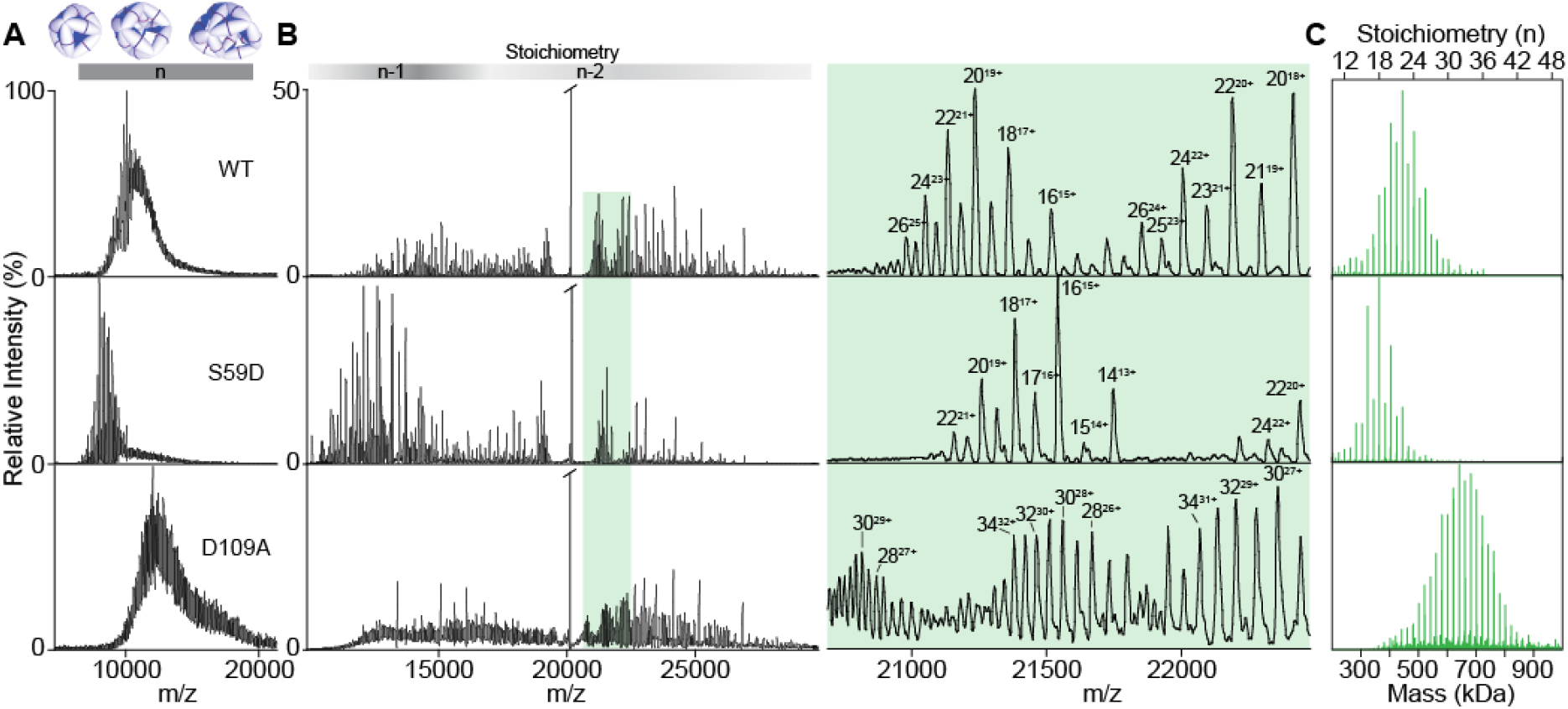
(A) Native MS of intact oligomeric assemblies of WT, Ser59Asp (phosphomimic), and Asp109Ala (isoAsp-mimic) αB, not directly assignable due to overlap of a multitude of charge states and stoichiometries. (B) Native MS with collision-induced dissociation (CID) of the assemblies in (A). Above shading shows regions where oligomers (n-mers) have lost one (n-1) or two (n-2) subunits. Detailed view of the region highlighted in green shows that with CID, charge states of unique oligomers are resolved. (C) Reconstructed oligomeric distributions. Data were charge deconvolved and then corrected to account for stripped subunits.

In addition to posing analytical challenges, isomerization and epimerization are also problematic from the perspective of molecular biology. It is not possible to use site-directed mutagenesis to insert D-residues or isoAsp into proteins, nor, given that both modifications are spontaneous, is it feasible to induce isomerization or epimerization in a site-specific and controllable fashion. In order to investigate the influence of isomerization at Asp109 on oligomeric assembly, we considered that our MD results had shown that the major consequence of modification from L-Asp was abrogation of the salt bridge with Arg120, and therefore mimicked this change by generating the mutant Asp109Ala. Native MS experiments on this mimic revealed it to be larger than wild-type, with a lower preference for even stoichiometries (Fig. 4C). To investigate whether this impact on the oligomeric distribution resulted in altered stability, we performed light-scattering experiments. The results show that Asp109Ala is significantly more prone to aggregation than wild type (Fig. S4). This is consistent with a similar tendency observed for the Arg120Gly variant, in which the same salt-bridge is disrupted.^21^ These results are consistent with a weakening of the dimer interface and the observation that the L-isoAsp variant is only observed in appreciable amounts in the insoluble fraction obtained from lenses (see Fig. 1f). Our isomerization mimicry therefore demonstrates that these changes in primary structure can impact oligomerization in the context of the full-length proteins.

## Conclusion

Isomerization and epimerization are prevalent PTMs in long-lived proteins such as the crystallins found in the eye lens. Although these modifications are difficult to detect and cannot be probed by site-directed mutagenesis, we have demonstrated that they can cause significant structural perturbation and loss of function. Epimerization of a single serine residue is sufficient to inhibit noncovalent recognition needed to maintain proper interface strengths and dynamics. Furthermore, epimerization of serine or other nearby residues alters phosphorylation and any functionality derived from it. By redirecting and altering the peptide backbone, aspartic acid isomerization can also inhibit kinase recognition and disrupt native salt bridge interactions, leading to improper oligomer assembly and size.

Previous studies have also noted structural changes brought about by similar modifications to proteins. For example, hen egg-white lysozyme with an L-isoAsp substitution at Asp101 causes a backbone deflection of nearly 90° relative to the native structure,^54^ and L-isoAsp32 insertion into ribonuclease forms a protruding U-shaped loop bent by nearly 90° instead of an α-helix.^55^ Isomerization can also affect physical properties such as solubility and bioactivity, with isomerization of Asp92 in immunoglobulin γ2 (IgG2) leading to deactivation of the antigen-binding region.^56^

These observations show that what might appear to be innocuous PTMs can disrupt protein structure. Given that long-lived proteins are also associated with many other diseases, including Alzheimer’s and Parkinson’s,^1,57,58^ it is likely that many of the structural issues highlighted herein also contribute to loss of function in these pathologies.

## Materials and Methods

### Lens samples

Human lenses were acquired from the National Disease Research Interchange (NDRI) (Philadelphia, Pennsylvania). The nuclei and cortices of the thawed lenses were separated using a 4mm trephine. The endcaps of each nucleus were removed by gently scraping off the outer tissue until only the dense nuclear portion remained. The nucleus and cortex were then separately homogenized in 600 μL of 50 mM Tris-HCl pH = 7.8. The lens fractions were then centrifuged at 15,100g for 20 min at 4ºC to separate the supernatant from the precipitate.

### WS fraction

The WS supernatant was purified by dialysis against water and lyophilized. 50 μg of lyophilized powder was added to 20 μL of 50 mM NH_4_HCO_3_ buffer, pH = 7.8. Disulfide bonds were then reduced with 1.5 μL of 100 mM of dithiothreitol (DTT) at 95°C for 5 minutes. After returning to room temperature, the solution was treated with 3 μL of 100 mM iodoacetamide (IAA) and incubated at room temperature in the dark for 20 minutes.

### WI fraction

The WI precipitate was solubilized in 6 M urea in 50 mM Tris-HCl pH=7.8 and purified by dialysis. Approximately 100 μg of the dialyzed protein (as determined by concentration based on absorbance at 280nm) was diluted to a total volume of 100 μL in 6 M urea in 50 mM Tris-HCl, pH = 8.0. Disulfide bonds were reduced with 5 μL of 200 mM DTT at 37°C for 20 minutes and then capped with 20 μL of 200 mM of IAA for one hour. The urea concentration was diluted to < 0.6 M using 50 mM Tris-HCl, 1 mM CaCl2, pH = 7.6.

Proteins were digested with trypsin for 12-16 hours at 37°C using a 50:1 protein to enzyme ratio. For samples requiring the iodobenzoate modification (i.e. RDD experiments), the digested peptides were desalted and cleaned using a peptide trap (Michrom Bioresource Inc). Approximately 5 nmoles of the digestion mixture, 10-100x excess of 4-iodobenzoic acid NHS-activated ester in dioxane and borate buffer (pH = 8.6) were combined and incubated for 1 hour at 37°C. Important: Dimethyl sulfoxide should not be substituted for dioxane in this step because it can cause aspartic acid isomerization. The modification side products at arginine and tyrosine side chains were removed by incubating the reaction mixture in 1 M hydroxylamine, pH = 8.5. The same procedure was used to modify synthetic peptide standards. These procedures have been determined previously not to yield any detectable isomerization in control experiments.^31^

### Peptide and Radical Precursor Synthesis

All synthetic peptides were synthesized manually using standard FMOC SPPS procedures with Wang resins, or Rink Amide resins as the solid support (29). N-hydroxysuccinimide (NHS) activated iodobenzoyl esters were synthesized by following a previous procedure. Synthetic model peptides were purified using a Jupiter Proteo column (250 mm x 4.6 mm, 4 μm, 90 Å, C12) (Phenomenex, Torrence, CA). These peptides were used to authenticate the identities of all peptide isomers and to generate calibration curves for accurate quantitation when full separation by LC was not achieved.

### Phosphorylation Assays

Mitogen-activated protein kinase-activated protein kinase 2 (MAPKAPK-2, also MK2) was purchased from Genetex (Irvine, CA). The kinase substrate peptides were synthesized with Rink Amide resins to prevent a charged C-terminus from interfering with kinase recognition. 100µM of each purified synthetic peptide was assayed in 25 mM HEPES buffer at pH 7.5 with 25mM MgCl_2_, and 2mM ATP in a 37 ºC sand bath. The assays were initiated by addition of 1µg kinase (for L-Ser and D-Ser assays) or 0.5µg kinase for (L-Asp, D-Asp, L-isoAsp, and D-isoAsp assays). Aliquots were taken at defined timepoints and the reaction was quenched by the addition of 1% TFA (v/v) before rapid freezing in liquid nitrogen. Samples were stored frozen until LCMS analysis. As a control, the all L- form of the FLRAPSWFDTG-NH_2_ peptide was assayed without the addition of kinase to eliminate the possibility of autophosphorylation.

### LC-MS Data Acquisition and Analysis

An Agilent 1100 series HPLC system (Agilent, Santa Clara, CA) with a BetaBasic column (150 mm x 2.1 mm, 3 μm, 150 Å, C18) was coupled to an LTQ mass spectrometer. Peptides were separated using a 0.1% formic acid in water (mobile phase A) and a 0.1% formic acid in acetonitrile (mobile phase B) binary system at a flow rate of 0.2 mL/min. The digestion mixtures were loaded onto the column and separated using the following gradient: 5% B to 20% B over 60 minutes, 20% B to 30% B over the next 45 minutes, 30% B to 50% B over the next 15 minutes and 50% B to 95% B over the final 10 minutes. The LTQ was operated in data-dependent mode using the Xcalibur program (Thermo Fisher Scientific). Specifically, in the CID-only LC-MS run, the first scan event was a full MS from m/z 400-2000 Da, followed by an ultrazoom (MS^2^) and then CID (MS^3^). In the RDD LC-MS experiments, the laser pulses were triggered during the MS^2^ step and CID was performed as a pseudo-MS^3^ step. Due to the high photodissociation yield of the 4-iodobenzoic acid chromophore, the major peak during this step is the loss of iodine, and it is the subsequent precursor for MS^3^. The exclusion time was set to 60 seconds for the identification of peptides. For isomer identification, an inclusion mass list was added and the exclusion time was reduced to 16 seconds to enable repeated analysis of isomers.

### R_isomer_ Calculations

To quantify isomer identification, the kinetic method using the R value, originally described by Tao and co-workers was utilized (31). In this paper, R_isomer_ represents the ratios of the relative intensities of a pair of fragments that varies the most between two isomers (R_A_/R_B_). Following acquisition of a tandem mass spectrum, R_isomer_ values are calculated for all pairs of peaks to reveal fragments that yield the best differentiation. If R_isomer_ = 1 then the two MS^n^ spectra are indistinguishable, and the species are likely not isomers. If R_isomer_ >1, a larger number indicates a higher probability that the two species are isomers. To confidently identify each of these isomers by MS^n^, we use a threshold that was determined by performing a *t*-test on the R_isomer_ values obtained by performing CID and RDD on a mixture of synthetic peptides separated by LCMS. Using 99% confidence intervals, the R_isomer_ threshold for CID is >1.9 and for RDD it is >2.4.^31^

### Protein expression and purification

Core αB (cABC, residues 68-153) was expressed in *E. coli* and purified as described previously.^47^ A gene insert encoding core αA (cAAC, residues 59-153) was purchased from Integrated DNA Technologies and inserted into a pET28a vector linearized with BamHI and XhoI (New England Biolabs) using an In-Fusion HD Cloning Kit (New England Biolabs) to generate a TEV-cleavable His-tagged construct. This was expressed and purified in the same manner as cABC with addition of 5 mM BME in all buffers prior to SEC, resulting in some population of BME-adducted protein visible in the spectra in Figure 2. Core domains were stored in 100 mM NaCl, 20 mM Tris, pH 8 at −80°C until use. Full length αB was expressed and purified as described previously.^47^ Mutations S59D and D109A were introduced using a Quik-Change Site-Directed Mutagenesis kit (Agilent) and mutants were expressed and purified in the same manner as WT. Full length proteins were stored in MS buffer (200 mM ammonium acetate pH 6.9) at −20°C until use. Concentrations were determined by UV absorbance at 280 nm.

### Native MS of core domains and peptides

Spectra were collected using a previously described protocol^59^ on a Synapt G1 IM-QToF mass spectrometer (Waters) with parameters as follows: capillary 1.5 kV, sampling cone 40 V, extraction cone 3 V, backing pressure 3.1 mbar, trap gas (argon) 3 mL min^−1^, trap cell voltage 10 V, transfer cell voltage 8 V. Ion mobility was enabled with parameters in the mobility cell: IMS gas flow 22 mL min^−1^, IMS wave velocity 320 m s^−1^, IMS wave height 5.5 V. Proteins were buffer exchanged into 200 mM ammonium acetate pH 6.9 using a Biospin-6 column (BioRad). All spectra were recorded at 10 μM based on concentration measurement post-buffer exchange. Samples were introduced using gold-coated capillaries prepared in-house. Lyophilized peptides -- Ct (ERAIPVSRE) and G-Ct-G (GERAIPVSRE) with L or D-Ser – were resuspended in milliQ H_2_O to a stock concentration of 1 mM and then diluted in MS buffer and mixed with protein immediately prior to analysis to a final concentration of 40 μM each for competition experiments. For quantitation, monomeric species were extracted in DriftScope (Waters) and intensities recorded from MassLynx using all resolved adduct peaks in addition to apo for 5+, 6+ and 7+ charge states. Data are reported as the mean +/- SD for three replicates.

### Native MS of full length αB and mutants

Spectra were collected on a modified QExactive hybrid quadrupole-Orbitrap mass spectrometer (ThermoFisher Scientific) optimized for transmission of high-mass complexes.^60^ Protein concentration was 15 μM by monomer. Capillary voltage was 1.4 kV in positive ion mode with source temperature 200°C and S-lens RF 200%. UHV pressure (argon) was between 1.4 − 10^−9^ and 1.7 − 10^−9^ mbar. In-source trapping fragmentation voltage ranged from −150 to −180 V. Ion transfer optics were as follows: injection flatapole 10 V, inter-flatapole lens 8 V, bent flatapole 6 V, transfer multipole 4 V, C-trap entrance lens 3 V. Nitrogen was used in the HCD cell and HCD energy was 0 V for intact spectra and tuned for optimal dissociation of each protein for CID spectra, ranging from 200 to 230 V. Resolution was kept at 17,500 at m/z = 200 for a transient time of 64 ms and the noise threshold was set to 3. For CID spectra, groupings of 30 microscans were combined to improve signal quality. Data were visualized using Xcalibur (ThermoFisher Scientific) and calibrated manually according to expected peak positions for WT αB-crystallin. Calibrated CID data were processed using UniDec software^61^ which allowed for stoichiometric assignment and post-hoc correction for dissociated subunits.

### Molecular modeling

PDB 4M5T, featuring cABC (residues 67-151) with each subunit complexed with a C-terminal peptide (156-164), was exploited to assess the interactions between C-terminal peptide and crystallin domains. Two molecular dynamics simulations were prepared, one with C-terminal peptides bound as shown in the crystal structure, i.e. running from the β8 to the β4 strands of cABC, and one with the peptide bound in the opposite direction, i.e. β4 to β8. In order to build a model of the β4-β8 peptide, we selected only the alpha carbons of β8-β4 peptide, renumbered them in the opposite direction, and used them as template to align a peptide extracted from the crystal structure. Structures were then simulated using the Amber ff14SB force field^62^ on the NAMD molecular dynamics engine^63^ Structures were first solvated in a box of TIP3P water, their box charge neutralized by addition of Na+ ions, and the resulting systems energy minimized with 2000 conjugate gradient steps. We then performed 0.5 ns steps in the NPT ensemble, with all protein’s alpha carbons constrained by a harmonic potential. Langevin dynamics were used to impose a temperature of 300 K, using a damping of 1 /ps. A constant pressure of 1 Atm was imposed via a Langevin piston having a period of 200 fs, and a decay of 50 fs. The system was then further equilibrated in the NVT ensemble for 1 ns, after which 200 ns production runs in the NPT ensemble were performed. In all simulation steps, Particle Mesh Ewald was used to treat long range electrostatic interactions, a cutoff distance of 12 Å was set on van der Waals interactions, and a 2 fs time step was exploited by restraining every covalent bond with SHAKE.

From each simulation, one frame per nanosecond was extracted from the production run for analysis. From each of these, hydrogen bonds between each residue part of a peptide and ones part of the crystallin domain were identified using VMD.^64^ We counted the occurrences of each bond in order to determine the percentage of time two residues spend interacting. A bond was considered established if the distance between donor and acceptor atoms was less than 3 Å, and the acceptor-donor-hydrogen angle less than 20 degrees. Residue pairs forming simultaneously more than one H-bond in the same frame would be counted as bonded only once.

In order to study the effect of Asp109 epimerization we exploited PDB 2WJ7, featuring a cABC dimer in Aβ II register (thus enabling the formation of the Asp109-Arg120 salt bridge). Three models featuring D-Asp, D-isoAsp and L-isoAsp at position 109 in one of the two monomers, respectively, were produced by modifying PDB 2WJ7 in Schrödinger Maestro. Simulation parameters for non-standard amino acids were produced with Antechamber^65,66^ All atom types could be assigned according to available ff14SB parameters. All models were simulated in explicit solvent for 150 ns following the same simulation protocol described above. His111-Arg123 and Asp109-Arg120 distances were measured every 100 ps. For the latter, we report the shortest distance between each of the hydrogens of Arg120 guanidinium, and oxygens of Asp109 carboxylate. The percentage of time a hydrogen bond is established was determined as described above, and averaged over the three simulations.

## Acknowledgments

The authors are grateful for funding from the NIH (NIGMS grant R01GM107099 to RRJ); Oxford University Press (Clarendon Award to MPC); Engineering and Physical Sciences Research Council (EP/P016499/1 to MTD, EP/J01835X/1 to JLPB). We thank Frances Kondrat and Gillian Hilton (University of Oxford) for initial mutagenesis work.

